# Dyslexia associated gene, *KIAA0319*, regulates cell cycle during human neuroepithelium development

**DOI:** 10.1101/2022.04.07.487504

**Authors:** Steven Paniagua, Bilal Cakir, Yue Hu, Ferdi Ridvan Kiral, Yoshiaki Tanaka, Yangfei Xiang, Benjamin Patterson, Jeffrey R. Gruen, In-Hyun Park

## Abstract

Reading Disability (RD), also known as dyslexia, is defined as difficulty processing written language in individuals with normal intellectual capacity and educational opportunity. The prevalence of RD is between 5% and 17%, and the heritability ranges from 44% to 75%. Genetic linkage analysis and genome-wide association studies (GWAS) have identified several genes and regulatory elements linked to RD and reading ability. However, their functions and molecular mechanisms are not well understood. Prominent among these is *KIAA0319*, encoded in the DYX2 locus of human chromosome 6p22. Association of *KIAA0319* has been independently replicated in multiple independent studies and languages. Rodent models suggest that *KIAA0319* is involved in neuronal migration, but its precise function is unknown. This studies aim to determine the mechanisms by which *KIAA0319* affects reading and language performance. We hypothesize that *KIAA0319* plays a critical role in neuronal development. RT-qPCR and quantitative immunofluorescence in the cortical neurons differentiated from H7 hESC show regulatory effects on proliferation and differentiation of neuronal progenitor cells. Knockdown of *KIAA0319* expression promotes early exit from the neuroepithelial cell stage and drives cells into cell cycle arrested neuronal progenitor cell stage. This suggests that *KIAA0319* act by regulating neurogenesis in the reading related centers of the brain by targeting the cell cycle of proliferative cells. This demonstrates how subtle changes in expression could affect an isolated trait such as reading without global brain effects.

## Introduction

Reading is a complex skill influenced by both environmental and genetic factors. Dyslexia, also known as reading disablilty, is characterized by reading difficulty in the setting of normal intelligence and an adequate education. In turn, individuals with Dyslexia generally experience persistent, long-term detrimental effects on educational achievement and socioeconomic status (Schatschneider and Torgesen, 2004). Dyslexia is the most common learning disability, with a prevalence of 5% to 17% and a heritability of 0.34 to 0.76 (DeFries et al., 1987; Hawke et al., 2006; Shaywitz and Shaywitz, 2005). Genetic studies have identified nine dyslexia loci (DYX1-9), contingent on aspects of reading performance(Fisher and DeFries, 2002; Gayan et al., 1999). The most frequently replicated locus is DYX2 on chromosome 6p21.3, and the most frequently replicated dyslexia genes on DYX2 are *KIAA0319* and *DCDC2* (Brkanac et al., 2007; Deffenbacher et al., 2004; Francks et al., 2004; Meng et al., 2005; Zhao et al., 2016).

Dyslexia genes are highly conserved. Animal models have provided insights into some of the cellular function of dyslexia genes (Paracchini et al., 2006; Szalkowski et al., 2012), However, as informative as animal models are they do not account for the complexity of reading or the critical skills that subserve reading such as decoding, phonological awareness, or orthographic coding, suggesting a need for modeling genetic variants associated with reading and dyslexia in human cells and tissues (Fitch, 2017; Kirby, 2017; Powers et al., 2016). While the role of *KIAA0319* in murine brain development has been defined, little is known about its role in human brain development. Previous animal studies showed that Kiaa319 interacts with intracellular trafficking protein AP-2. Additionally, Kiaa0319 undergoes N- and O-glycosylation, mostly associated with the plasma membrane. Both studies suggest that Kiaa0319 plays a role in cell maintenance and cell-cell signaling pathway in the endocytosis pathway (Levecque et al., 2009; Velayos-Baeza et al., 2008). Kiaa0319 has also been found to be associated with extracellular signaling pathways (Franquinho et al., 2017; Wu et al., 2020), via Smad2/3, which affects global gene expression. Both endocytosis and extracellular signaling pathways are essential for regulating neurogenesis (Cope and Gould, 2019).

Functional studies of dyslexia genes suggested that they are involved in signaling and radial migration of terminally differentiated neurons were the prime focus of dyslexia gene function in the brain (Che et al., 2016). However, it is neuronal epithelia cells and neuronal progenitor cells stages of differentiation, well before terminal differentiation, that radial migrate and populate individual cortical layers (Cope and Gould, 2019). Neuroepithelial cells perform radial migration and populate the individual cortical layers of the brain as proliferative neuronal progenitor cells (Ji et al., 2017; Stockinger et al., 2011). Neuronal progenitor cells act as a critical inflection point between neurons, astrocytes, and oligodendrocytes, providing the diversity in neuronal subtypes required essential for neural circuitry’s construction, maintenance, and efficiency for brain function, particularly higher brain function such as reading (Bergles and Richardson, 2015; Durkee and Araque, 2019; Gotz and Huttner, 2005; Klausberger and Somogyi, 2008; Molofsky and Deneen, 2015). Additionally, neuronal progenitor cells regulation is crucial for neurogenesis within the adult brain, which has been associated with learning and memory by propagating new adult neurons stemming from the neuronal progenitor cell population housed within the dentate gyrus (Cameron and Glover, 2015).

Over the past 10 years, gene editing in pluripotent cells such as human embryonic stem cells (hESCs) and induced pluripotent stem cells (iPSCs) have been shown to be an informative approach to modeling the effects of genetic variants on neuronal differentiation. Pluripotent stem cell stems can be induced to differentiate into many cell types, including glutamatergic neurons, GABAergic neurons, and glial cells that populate the human brain (Damdimopoulou et al., 2016). Neuroectodermal differentiation of hESCs can reproduce embryonic brain development stages and gene expression patterns (Howard et al., 2008). Combining with the gene editing tools, hESCs provide powerful experimental tools to dissect the function of genes associated with neurodevelopmental disorders or neuropsychiatric disorders (Kolagar et al., 2020).

We the hypothesize that the clinical association of *KIAA0319* with reading performance and dyslexia is a consequence of altered expression on neurogenesis during early stages of embryonic brain development. To test this hypothesis, we modeled *KIAA0319* expression knockdown in hESCs induced into neuroectodermal differentiation and measured the effects on transitions through stages of typical neuronal development. We found that KIAA0319 is essential for neuroepithelial cell differentiation, which could affect radial migration as well as downstream differentiation into diverse populations of brain cells, providing clues as to how this gene could influence complex traits such as reading.

## Results

### *KIAA0319* regulates the transition of neuronal epithelial cells into neuronal progenitor cells

To examine *KIAA0319* gene expression pattersn in human, we analyzed transcriptome profiles of human somatice tissues and during human fetal development. KIAA0319 is more strongly expressed throughout human cortical development and in all brain regions (**Figure S1A**). Throughout human fetal development *KIAA0319* expresssion in brain is elevated and is then maintained after birth (**Figure S1B-D**, (Cardoso-Moreira et al., 2019; Pletikos et al., 2014)). Importantly, in human cortinal neural differentiation of hESCs, we found that KIAA0319 showed similiar expression pattern like cortical fetal development (**Figure S1E**), suggesting that our hESC-derived cortical neurons reflect similar molecular patterns with *in vivo* human fetal development.

In order to study the function of KIAA0319, we use CRISPRi based on the dCas9-KRAB to deplete expression of KIAA0319 in hESCs (**Figure 1A**). Both qRT-PCR and immunofluorescence confirmed the depletion of expression of *KIAA0319* in CRISPRi cells compared to controls during neuronal differentiation at days 7, 14, 21, and 28 (**Figure 1B-D**). To examine the effects of *KIAA0319* knockdown on differentiation, we surveyed the expression of gene markers emblematic of transtional stages of neuronal differentiation genes by qRT-PCR and immunostaining on days 7, 14, 21, and 28 of cortical differentiation. We focused on crucial neuronal stage lineage markers *SOX10, PAX6, TBR2, TBR1*, and *MAP2* to quantitate the population of neuroepithelial, neuronal progenitor, immature neuron, and mature neurons, respectively. Compared to controls SOX10 expression by qRT-PCR and immunofluorescense, a marker of neuronal epithelial cells, was suppressed at day 21 in *KIAA0319* KD cells (**Figure 2A-C and E**). Conversely, PAX6 expression by qRT-PCR and immunofluorescence, a marker of neuronal progenitor cells, increased in *KIAA0319* KD cells on day 14 and 21 (**Figure 2D-F**). Together, the results of these experiments suggest that *KIAA0319* is critical for regulating the transition of SOX10+ neuronal epithelial cells to PAX6+ neuronal progenitor cells.

**Figure 1.**
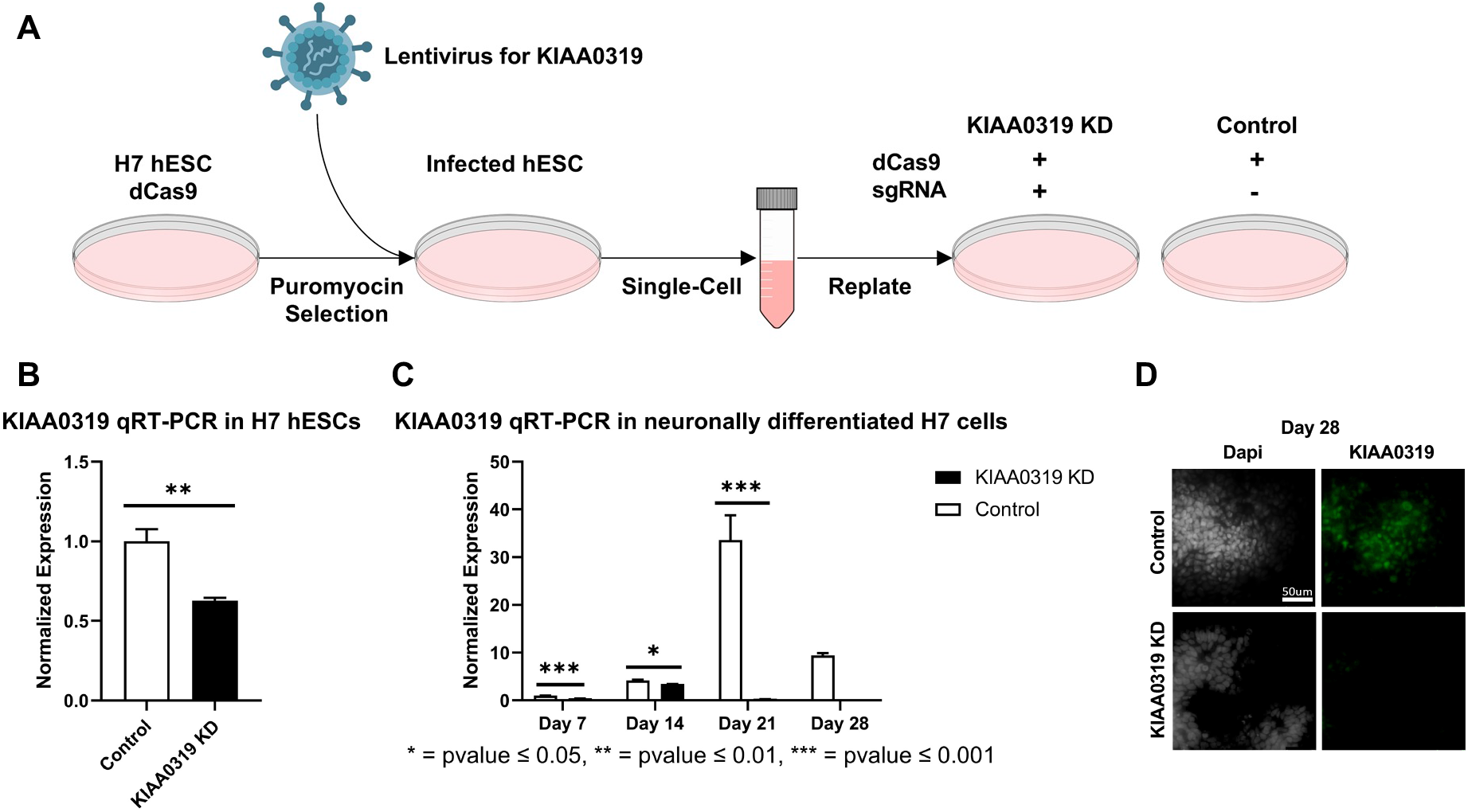
Depletion of *KIAA0319* by CRISPR interference (CRISPRi) **(A)** A diagram to deplete the expression of KIAA0319 by using CRISPRi in hESCs. **(B)** Confirmation of the depletion of *KIAA0319* by qRT-PCR at H7 cells after transfection and puromycin selection. (**C**) Confirmation of the supression of *KIAA0319* by qRT-PCR on days 7, 14, 21, and 28 of neuronal differentiation. (**D**) Immunofluorescence images of *KIAA0319* KD and Control on day 28. Green: KIAA0319, grey: DAPI. Scale bar: 50 µm.

**Figure 2.**
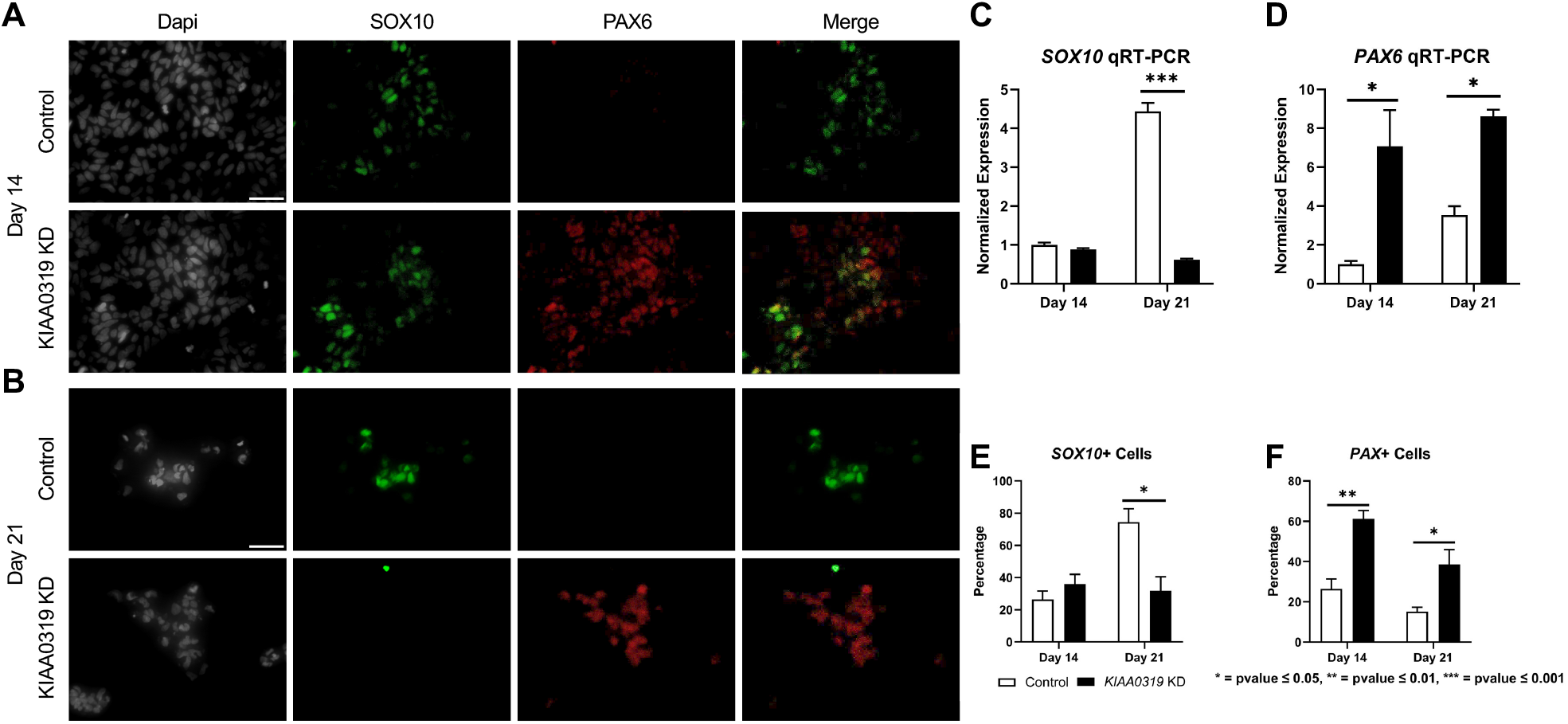
Depletion of *KIAA0319* decreases expression of neuronal epithelial marker SOX10 but increases neuronal progenitor marker PAX6. (**A**) After 14 days of neuronal differentiation, immunofluorescence was performed for SOX10 (green), PAX6 (red), and DAPI (grey). (**B**) After 21 days of neuronal differentiation, immunofluorescence was performed for SOX10 (green), PAX6 (red), and DAPI (grey). qRT-PCR for SOX10 (**C**) and PAX 6 (**D**) were performed on days 14 and 21 of *KIAA0319* KD and Control H7 cells. The percentage of the population of SOX10 (**E**) or PAX6 (**F**) positive cells were calculated on day 14 and day 21 of differentiation for *KIAA0319* KD and Control H7 cells.

### *KIAA0319* KD arrests cells in a non-proliferative neuronal progenitor cell stage

We further characterized how *KIAA0319* regulates PAX6 positive neural progenitor cells at later stages of neuronal differentiation. We simultaneously quantified KI67, a marker for proliferation, alongside PAX6 by qRT-PCR and immunofluorescence to quantify the proliferation of neuronal progenitor cells at two late time points of differentiation (day 28 and 42) (**Figure 3A-B**). We found that the fraction of KI67+ cells remained relatively low in *KIAA0319* KD cells, compared to control (**Figure 3C**), while the fraction of PAX6+ cells was relatively the same (**Figure 3D**). However, the fraction of KI67+ cells among the PAX6+ cells was decreased, suggesting an overall decrease in the population of proliferative neuronal progenitor cells (**Figure 3E**). Overall, these results suggest that KIAA0319 knockdown is either driving the PAX6+ neuronal progenitor cells into cell cycle arrest, inducing apoptosis in the proliferate population, or prematurely transitioned into downstream stages of neuronal differentiation.

**Figure 3.**
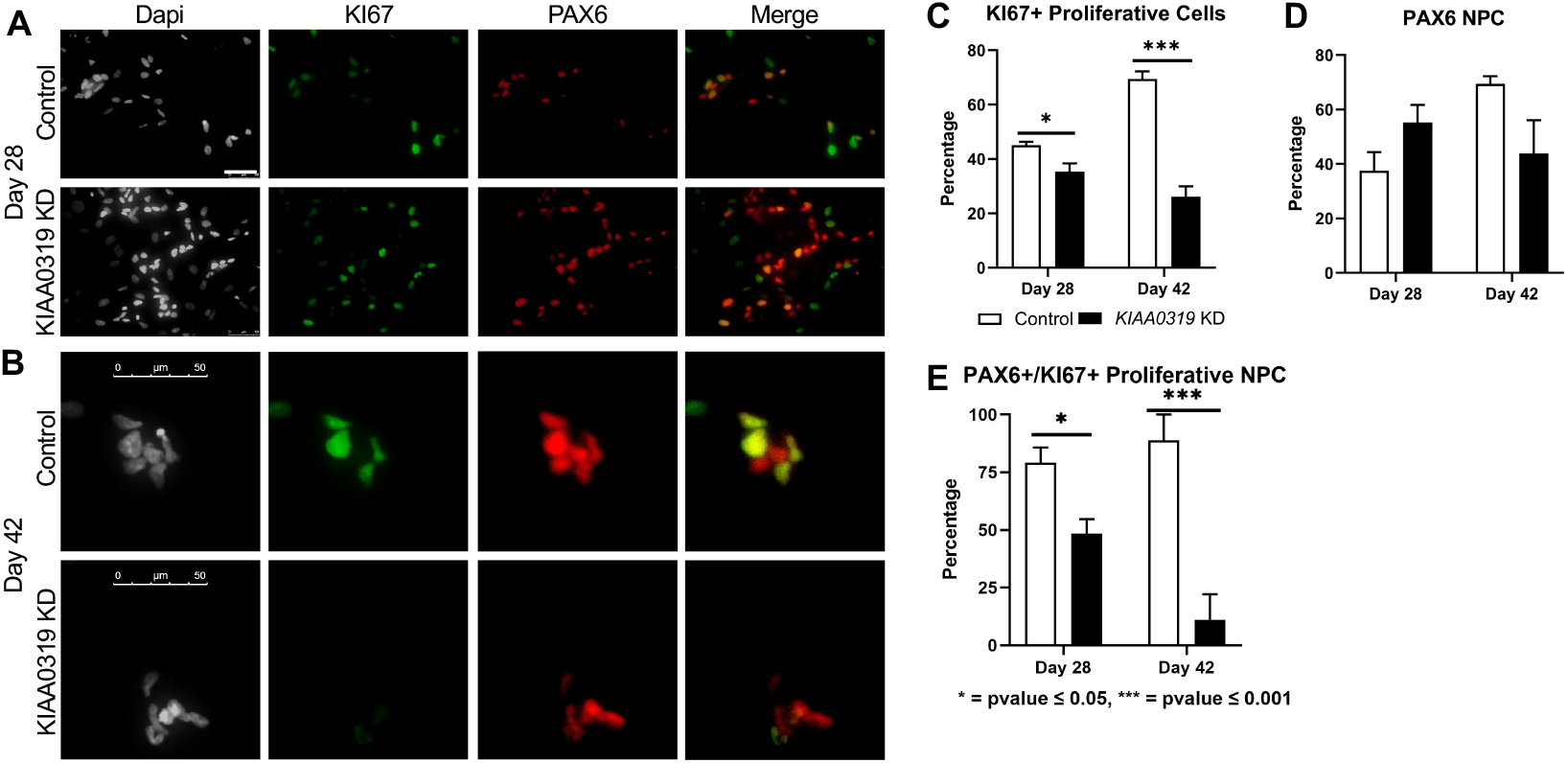
Depletion of *KIAA0319* decreases the percentage of proliferative neuronal progenitor cells. **(A-B)** Immunostaining of KIAA0319 KD and control cells for KI67 (green) and PAX6 (red) in Dapi (gray) on day 28 (**A**) and day 42 (**B**) of neuronal differentiation. Scale bar: 50 µm. Percentage of KI67+ cells **(C)** and PAX6 positive population **(D)** among Dapi positive cells at day 28 and day 42. Percentage of PAX6 positive cells among KI67 positive population at day 28 and day 42 (**E**).

To assess whether the decrease in KI67+ PAX+ neuronal progenitor cells was not due to apoptosis, we stained knockdown and control cells with cleaved caspase-3 antibody, a marker of cell death induction at day 28 (Figure 4A). *KIAA0319* KD did not induce apoptosis suggesting that *KIAA0319* that the decrease in KI67+ PAX6+ neuronal progenitor cells was not due to apoptosis. Following this, we examined markers at later stages of neuronal differentiation to quantify if *KIAA0319* knockdown induces premature transition from neuronal progenitor cells into downstream stage of neuronal differentiation. Compared to controls at day 28, *KIAA0319* knockdown significantly decreased expression of markers of intermediate progenitor cells (TBR2), immature neuronal cells (TBR1 and TUJ1), and mature neuronal cells (MAP2) (**Figure 4B**). The results of these experiments show that the knockdown arrests cells in a non-proliferative neuronal progenitor cell stage, and suggest that *KIAA0319* is important for transitioning from neuronal epithelial cells through neuronal progenitor cells and downstream stages of neuronal differentiation.

**Figure 4.**
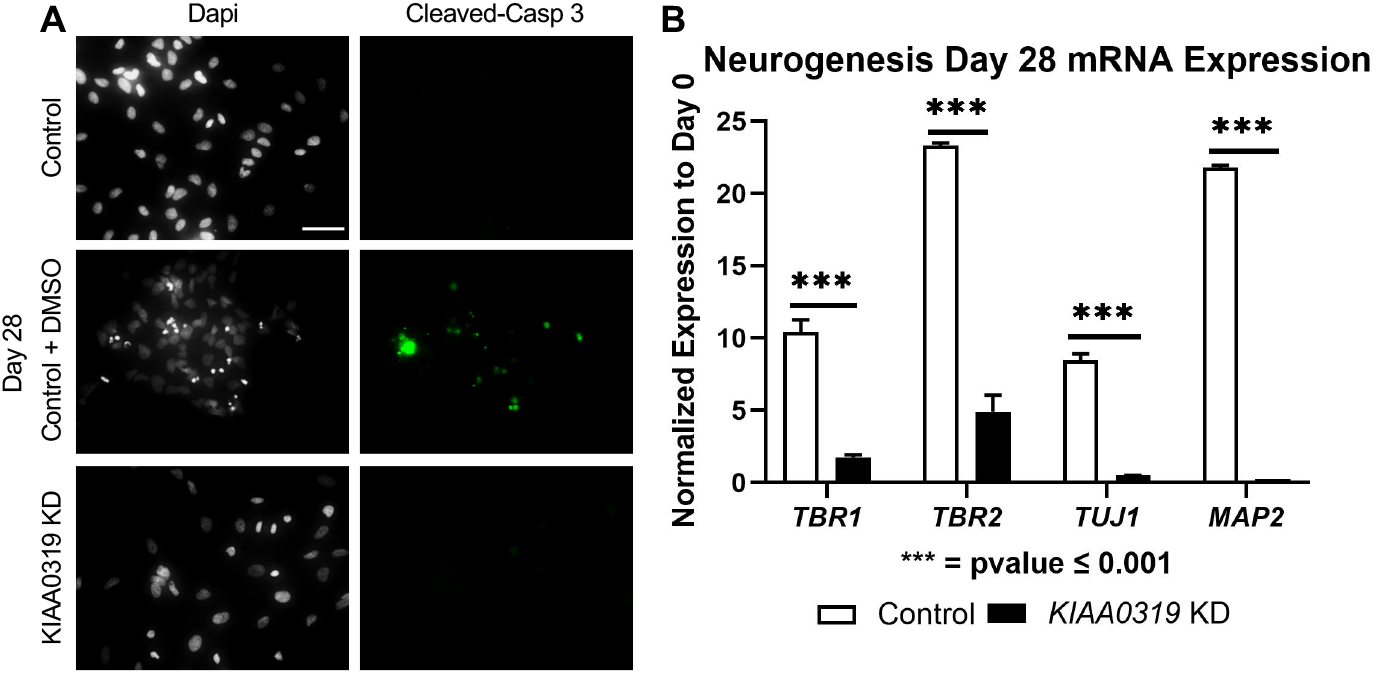
*KIAA0319* KD cells do not continue differentiating. (**A**) Images of *KIAA0319* KD, Control, Control + DMSO (Positive Apoptosis Control) at day 28 of neuronal differentiation stained with cleaved-caspase 3 (green) and Dapi (grey). (**B**) QRT-PCR at day 28 for neuronal lineage markers *TBR1, TBR2*, and *MAP2*.

### *KIAA0319* KD drives neuronal progenitor cells into cell cycle arrest

To examine how *KIAA0319* could be regulating proliferation in neuronal progenitor cells, we performed RNA-sequencing on the same aliquots of cells as the experiments about at day 7, 14, 21, and 28 (**Figure 5A**). We identified 85-257 differentially expressed genes between *KIAA0319* KD and control at each development stage **(Figure S2-5)**. Ingenuity Pathway Analysis (IPA) to differentially expressed genes revealed that pathways of endocytosis, exocytosis, lysosome, differentiation, proliferation, and metabolism are dysregulated in *KIAA0319* KD (**Figure 5B**). Interestingly, all of these pathways have been previously implicated in driving cell cycle arrest in neuronal progenitor cells (Harada et al., 2021; Kobayashi et al., 2019; Zhang et al., 2021). Additionally, the statistically significant differentially expressed genes show a consistent decrease in differentiation associated genes at neuroepithelial and neuronal progenitor cell stages (**Figure 5C**). The changes in the extracellular matrix, metabolism, cell maintenance, and cell-cell signaling further support the findings that *KIAA0319* KD drives neuronal progenitors into cell cycle arrest.

**Figure 5.**
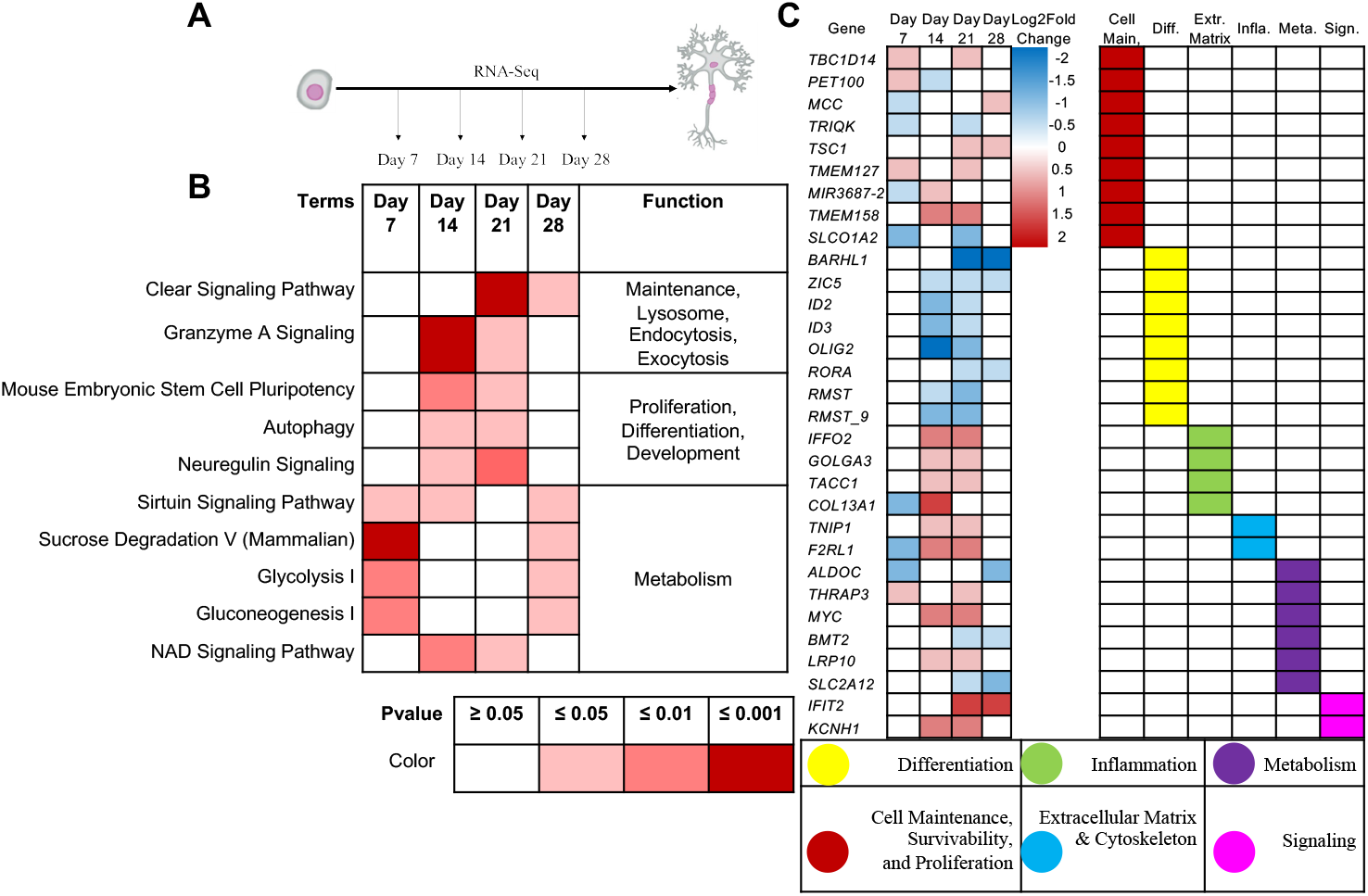
*KIAA0319* KD RNA-Seq shows pathways that promote cell cycle arrest of neuronal progenitor cells. (**A**) RNA Sequencing is performed at four distinct timepoints separated by seven-day intervals during cortical neuronal differentiation. (**B**) Ingenuity Pathway Analysis revealed enrichment of pathways appearing in multiple time points. (**C**) Ingenuity Pathway Analysis revealed enrichment of genes appearing in multiple time points.

## Discussion

In this study we set out to test our hypothesis that the clinical association of *KIAA0319* with reading performance and dyslexia is a consequence of altered expression on neurogenesis during early stages of embryonic brain development. We modeled knockdown of *KIAA0319* expression in hESCs induced into neuroectodermal differentiation and measured the effects on transitions through stages of typical neuronal development. *KIAA0319* depletion showed a decrease in the SOX10 neuroepithelial cell marker concurrent with an increase in the PAX6 neuronal progenitor marker at early stages of neurogenesis, and suppression of downstream stages of neurogenesis. These findings are consistent with a decreased proliferative state due to cell cycle arrest at the neuronal progenitor cell stage, highlighting the importance of tight regulation of *KIAA0319* expression during a critical period of brain development.

Low expression of *KIAA0319* appears to promote early exit from the neuroepithelial cell stage and entry into the neuronal progenitor cell stage. This suggests that KIAA0319 plays a critical role as an upstream gatekeeper for neurogenesis, slowing or speeding downstream differentiation by arresting cells at the neuronal progenitor cell stage. Transcriptome analysis suggests that *KIAA0319* could be regulating cell proliferation through endocytosis-related pathways. Dysregulation of the endocytosis pathways has been shown to drive neuronal progenitor cells into a stage of cell cycle arrest (Zhang *et al*., 2021). Our results provide a new avenue for studying the regulation of neurogenesis in human neurons.

This is the first study to suggest that *KIAA0319*, which has been associated with reading performance in multiple independent studies, may act by regulating neurogenesis. The results suggest that *KIAA0319* expression is required during the early stages of neurogenesis, centered around controlling the proliferation and differentiation of neuronal progenitor cells. The population of mature adult neurons is tightly controlled by the sustained proliferative populations of active neuronal progenitors and, while knockdown drives neuronal progenitors into a state of cell cycle arrest, this suggests *KIAA0319* plays a role in this highly regulated process. The *KIAA0319* genetic variants associated with reading and dyslexia are located in the 5-prime promoter region and regulate expression. This is consistent with a tightly regulated process whereby subtle changes in expression could affect an isolated high-order trait such as reading without global brain effects, typical of children and adults with dyslexia who by definition have normal or above normal intellectual ability.

## Methods

### Cell Culture

HEK 293T cells were maintained in 6-well plates at 37°C with 5%CO_2_. Cells were tested for mycoplasma and passaged every 3 days using Gibco TrypLE™ Express with Phenol Red for dissociation. Culture media consisted of 88% v/v DMEM, 10% v/v FBS, 1% v/v Glutamax, and 1% v/v Penicillin/Streptomycin. H7^dCas9-KRAB-b6^ (WiCell, WA07, H7) human ESCs were maintained in feeder-free culture conditions. Using Matrigel-coated cell culture dishes and mTeSR1 media at 37°C and 5%CO_2_. Cells were tested for mycoplasma and passaged every 7 days following dissociation with Dispase (0.83 U/ml). All experiments involving hESCs were approved by the Yale Embryonic Stem Cell Research Oversight Committee (ESCRO).

### Neuronal Differentiation

Pre-differentiated cells were removed before hESC colonies, dissociated into single cells using Accutase, and plated on 24-well plates at 500,000 cells/well with 50 µM Y27632 and mTeSR1. Once cultures reached 90% confluency, cells were grown for 10 days with neural induction media (47.5% v/v; DMEM/F12, 47.5% v/v Neurobasal media, 2% v/v B27 supplement, 1% v/v DMEM-NEAA, 1% v/v N2 Supplement, 1% v/v Glutamax, 20 µg/mL Insulin, 100 nM LDN-193189, 50 µM β-Mercaptoethanol, 10 µM SB431542, 2 µM XAV939), changing media every day. Cells were dissociated using Accutase on day 11 and plated on new 24-well plates along with 50 µM Y27632 and progenitor expansion media (neural induction media without inhibitors: LDN-193189, SB431542, and XAV939), changing media every other day. On day 19, culture media was changed to maturation media (95% v/v Neurobasal media, 2% v/v B27 supplement, 1% v/v N2 Supplement, 1% v/v Glutamax, 1% v/v Penicillin/Streptomycin) with 25ng/mL BDNF, changing media every two days for two weeks and every four days thereafter. Cells were dissociated using Accutase and replated on new 24-well plate on day 25 at 100,000 cells/well when confluency was reached.

### *KIAA0319* Lentivirus

Single Guide RNA (sgRNA) and reverse complement were designed using CRISPR PAM sites (crispr.mit.edu) targeting the transcription start site of *KIAA0319* (*KIAA0319* KD plasmid) and synthesized by Keck (sgRNA *KIAA0319* KD and Control listed in Table 2). Oligos were annealed using 50% v/v NEB Annealing Buffer, 46% v/v ddH_2_0, 2% v/v top oligo, and 2% v/v bottom oligo (Oligos listed in Table 2). Oligos were incubated at 95°C and left to cool to room temperature. Plasmid pCRISPia-V2, encoding a Puromycin resistance gene, had a sgRNA insertion site opened using digestion enzymes BstXI and Blpl. Ligation reactions consisted of 70% v/v ddH_2_O, 10% v/v of 10X T4 ligase buffer, 10% v/v 1:20 diluted annealed oligos, 5% pCRISPia-V2 (totaling 100ng of digested plasmid), and 5% v/v T4 Ligase at room temperature for 1 hour.

**Table 1:**
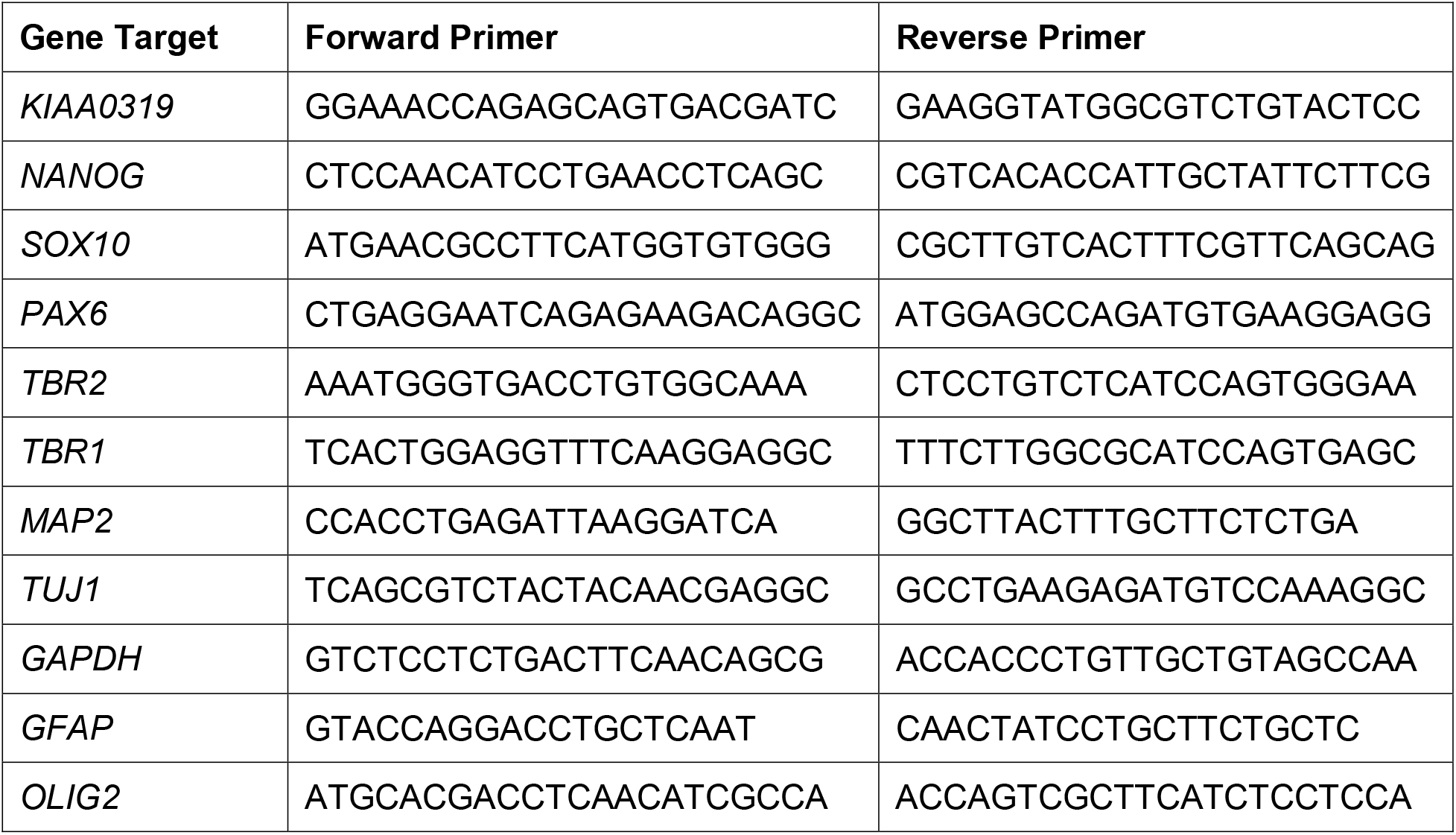
qRT-PCR gene target primers

**Table 2:**
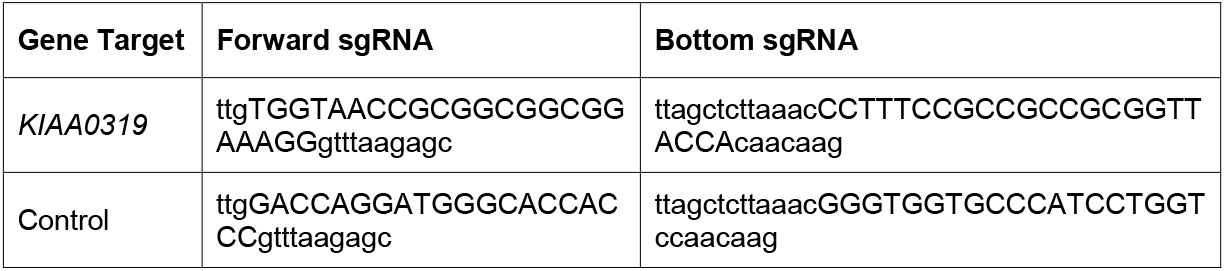
CRISPRi Oligos

Ligation reactions were transformed into One Shot™ TOP10 Chemically Competent cells following manufactures instructions and plated on 10cm LB Agar + Ampicillin(100ug/ml) plates from Recombinant Technologies, LLC at 37°C for culture overnight. Surviving colonies were cultured overnight in Gibco LB Broth with Ampicillin at 100ug/ml in a 37°C shaker, followed by plasmid purification using QIAGEN Plasmid Mini Kit and sequence confirmation.

Previously mentioned *KIAA0319* KD and Control plasmids were individually transfected into HEK293T cells along with 2.5 ugs of packaging plasmids using X-treme GENE 9 DNA transfection reagent following manufacturer’s instructions. Supernatants containing lentivirus were collected and concentrated 48 hours after transfection and stored at −80°C. H7 cells were infected with *KIAA0319* KD lentivirus and Control lentivirus. Infected cultures were selected following puromycin selection for 5 days.

### Real Time Quantitative PCR

Four wells of a 24-well plate of H7 cells transfected with *KIAA0319* KD or control samples were collected on days 7, 14, 21, and 28 of neuronal differentiation per replicate. Samples were homogenized using QIAGEN QIAshredder columns, and total RNA was isolated using the RNeasy Mini Kit following the manufacturer’s instructions. 1 ug of total RNA was used to synthesize cDNA using amfiRivert cDNA Synthesis Platinum Master Mix. Real-time quantitative PCR (RT-PCR) was performed using amfiSure qGreen Q-PCR Master Mix (2X), Low Rox on a CFX96 R-PCR System at amfiSure manufacture cycling conditions. Gene expression was quantified using the ΔΔC_T_ method with *GAPDH* as a housekeeping gene control.

### Immunostaining

Twelve wells of a 24-well plate of *KIAA0319* KD or Control cells were washed once in PBS before fixing in 4% Formaldehyde/PBS at room temperature for 15 minutes and then washed in PBS (3 x for 15 mins). In addition, samples were permeabilized by incubating with 0.1% Triton-100/PBS at room temperature for one hour and then washed in PBS (3 x for 15 mins) before storing at 4°C in PBS.

Blocking was done with 3% BSA/0.1% Triton-100/PBS at 4°C for two hours before incubating with primary antibodies diluted to the manufacturers recommendation in 3% BSA/0.1% Triton-100/PBS at 4°C overnight. Samples were then washed in PBS (3 x for 15 mins) before incubating with secondary antibody diluted in 3% BSA/0.1% Triton-100/PBS at room temperature for 1 hour. Samples were washed in PBS (3 x for 15 mins) before staining with DAPI to highlight nuclei for five minutes before final PBS wash. Immunofluorescence images were captured on a Leica TCS SP5 Spectral Confocal Microscope using Leica LAS AF software. Images were processed using ImageJ-Fiji.

### RNA-Sequencing

Cell pellets for three biological replicates of *KIAA0319* KD and control for days 7, 14, 21, and 28 for collected and frozen. Cell pellets were then processed to isolated RNA for library prep, and NOVA-seq for 50 mil reads per sample with a pair-end 150 bp at Yale Center for Genomic Analysis (YCGA).

FASTQ files were processed using an RNA-Sequencing pipeline developed from YCGA. Raw RNA-Sequencing reads were trimmed and aligned using Hierarchical Indexing for Spliced Alignment of Transcripts 2 (HISAT2)(Kim et al., 2019). Aligned reads are then concatenated using StringTie/Ballgown, followed by quality control using Picard(Frazee et al., 2015; Pertea et al., 2015). Finally, R package DESeq2 was used to generate summary reports and heatmaps using default setting (Love et al., 2014) (**Supplemental Figure 2-5**).

### Ingenuity Pathway Analysis (QIAGEN IPA)

https://digitalinsights.qiagen.com/products-overview/discovery-insights-portfolio/analysis-and-visualization/qiagen-ipa/

IPA provided core analysis based on DESeq2 selected significant genes to identify significantly changed pathways. Filtered for pathways and genes present at multiple timepoints and then grouped by GO Term.

## Figure Legends

**Supplemental Figure 1:** (**A**) Bult Tissue Expression for *KIAA0319* provided by GTEx. Transcripts per million from all sampled brain regions in the GTEx project. (**B**) Bulk RNA-Sequencing provided by (Cardoso-Moreira *et al*., 2019) and selecting for *KIAA0319* expression. Expression data provided by (Pletikos *et al*., 2014) available from the Human Brain Transcriptome Database, (**C**) KIAA0319 signal intensity within the Neonatal Cortex. (**D**) *KIAA0319* signal intensity within the Brain Regions. cerebellar cortex (CBC), mediodorsal nucleus of the thalamus (MD), striatum (STR), amygdala (AMY), hippocampus (HIP) and 11 areas of neocortex (NCX): Frontal cortex (OFC, DFC, VFC, MFC, M1C), Parietal cortex (S1C, IPC), Temporal cortex (A1C, STC, ITC), Occipital cortex (V1C).

**Supplemental Figure 2: Significant genes between KIAA0319 Knockdown and Control at day 7**. (**A**) Heatmap for all 99 significant genes (**B**) Volcano plot for all significant genes determined by adjusted pvalue (**C**) Boxplot for top 30 significant genes. (**D**) Individual Boxplots for the top 30 significant genes.

**Supplemental Figure 3: Significant genes between KIAA0319 Knockdown and Control at day 14**. (**A**) Heatmap for all 85 significant genes (**B**) Volcano plot for all significant genes determined by adjusted pvalue (**C**) Boxplot for top 30 significant genes. (**D**) Individual Boxplots for the top 30 significant genes.

**Supplemental Figure 4: Significant genes between KIAA0319 Knockdown and Control at day 21**. (**A**) Heatmap for all 257 significant genes (**B**) Volcano plot for all significant genes determined by adjusted pvalue (**C**) Boxplot for top 30 significant genes. (**D**) Individual Boxplots for the top 30 significant genes.

**Supplemental Figure 5: Significant genes between KIAA0319 Knockdown and Control at day 28**. (**A**) Heatmap for all 89 significant genes (**B**) Volcano plot for all significant genes determined by adjusted pvalue (**C**) Boxplot for top 30 significant genes. (**D**) Individual Boxplots for the top 30 significant genes.

## Notes

### Competing Interest Statement

The authors have declared no competing interest.

## References

Bergles, D.E., and Richardson, W.D. (2015). Oligodendrocyte Development and Plasticity. Cold Spring Harb Perspect Biol 8, a020453. 10.1101/cshperspect.a020453.

Brkanac, Z., Chapman, N.H., Matsushita, M.M., Chun, L., Nielsen, K., Cochrane, E., Berninger, V.W., Wijsman, E.M., and Raskind, W.H. (2007). Evaluation of candidate genes for DYX1 and DYX2 in families with dyslexia. Am J Med Genet B Neuropsychiatr Genet 144B, 556–560. 10.1002/ajmg.b.30471.

Cameron, H.A., and Glover, L.R. (2015). Adult neurogenesis: beyond learning and memory. Annu Rev Psychol 66, 53–81. 10.1146/annurev-psych-010814-015006.

Cardoso-Moreira, M., Halbert, J., Valloton, D., Velten, B., Chen, C., Shao, Y., Liechti, A., Ascencao, K., Rummel, C., Ovchinnikova, S., et al. (2019). Gene expression across mammalian organ development. Nature 571, 505–509. 10.1038/s41586-019-1338-5.

Che, A., Truong, D.T., Fitch, R.H., and LoTurco, J.J. (2016). Mutation of the Dyslexia-Associated Gene Dcdc2 Enhances Glutamatergic Synaptic Transmission Between Layer 4 Neurons in Mouse Neocortex. Cereb Cortex 26, 3705–3718. 10.1093/cercor/bhv168.

Cope, E.C., and Gould, E. (2019). Adult Neurogenesis, Glia, and the Extracellular Matrix. Cell Stem Cell 24, 690–705. 10.1016/j.stem.2019.03.023.

Damdimopoulou, P., Rodin, S., Stenfelt, S., Antonsson, L., Tryggvason, K., and Hovatta, O. (2016). Human embryonic stem cells. Best Pract Res Clin Obstet Gynaecol 31, 2–12. 10.1016/j.bpobgyn.2015.08.010.

Deffenbacher, K.E., Kenyon, J.B., Hoover, D.M., Olson, R.K., Pennington, B.F., DeFries, J.C., and Smith, S.D. (2004). Refinement of the 6p21.3 quantitative trait locus influencing dyslexia: linkage and association analyses. Hum Genet 115, 128–138. 10.1007/s00439-004-1126-6.

DeFries, J.C., Fulker, D.W., and LaBuda, M.C. (1987). Evidence for a genetic aetiology in reading disability of twins. Nature 329, 537–539. 10.1038/329537a0.

Durkee, C.A., and Araque, A. (2019). Diversity and Specificity of Astrocyte-neuron Communication. Neuroscience 396, 73–78. 10.1016/j.neuroscience.2018.11.010.

Fisher, S.E., and DeFries, J.C. (2002). Developmental dyslexia: genetic dissection of a complex cognitive trait. Nat Rev Neurosci 3, 767–780. 10.1038/nrn936.

Fitch, W.T. (2017). Empirical approaches to the study of language evolution. Psychon Bull Rev 24, 3–33. 10.3758/s13423-017-1236-5.

Francks, C., Paracchini, S., Smith, S.D., Richardson, A.J., Scerri, T.S., Cardon, L.R., Marlow, A.J., MacPhie, I.L., Walter, J., Pennington, B.F., et al. (2004). A 77-kilobase region of chromosome 6p22.2 is associated with dyslexia in families from the United Kingdom and from the United States. Am J Hum Genet 75, 1046–1058. 10.1086/426404.

Franquinho, F., Nogueira-Rodrigues, J., Duarte, J.M., Esteves, S.S., Carter-Su, C., Monaco, A.P., Molnar, Z., Velayos-Baeza, A., Brites, P., and Sousa, M.M. (2017). The Dyslexia-susceptibility Protein KIAA0319 Inhibits Axon Growth Through Smad2 Signaling. Cereb Cortex 27, 1732–1747. 10.1093/cercor/bhx023.

Frazee, A.C., Pertea, G., Jaffe, A.E., Langmead, B., Salzberg, S.L., and Leek, J.T. (2015). Ballgown bridges the gap between transcriptome assembly and expression analysis. Nat Biotechnol 33, 243–246. 10.1038/nbt.3172.

Gayan, J., Smith, S.D., Cherny, S.S., Cardon, L.R., Fulker, D.W., Brower, A.M., Olson, R.K., Pennington, B.F., and DeFries, J.C. (1999). Quantitative-trait locus for specific language and reading deficits on chromosome 6p. Am J Hum Genet 64, 157–164. 10.1086/302191.

Gotz, M., and Huttner, W.B. (2005). The cell biology of neurogenesis. Nat Rev Mol Cell Biol 6, 777–788. 10.1038/nrm1739.

Harada, Y., Yamada, M., Imayoshi, I., Kageyama, R., Suzuki, Y., Kuniya, T., Furutachi, S., Kawaguchi, D., and Gotoh, Y. (2021). Cell cycle arrest determines adult neural stem cell ontogeny by an embryonic Notch-nonoscillatory Hey1 module. Nat Commun 12, 6562. 10.1038/s41467-021-26605-0.

Hawke, J.L., Wadsworth, S.J., and DeFries, J.C. (2006). Genetic influences on reading difficulties in boys and girls: the Colorado twin study. Dyslexia 12, 21–29. 10.1002/dys.301.

Howard, B.M., Zhicheng, M., Filipovic, R., Moore, A.R., Antic, S.D., and Zecevic, N. (2008). Radial glia cells in the developing human brain. Neuroscientist 14, 459–473. 10.1177/1073858407313512.

Ji, L., Bishayee, K., Sadra, A., Choi, S., Choi, W., Moon, S., Jho, E.H., and Huh, S.O. (2017). Defective neuronal migration and inhibition of bipolar to multipolar transition of migrating neural cells by Mesoderm-Specific Transcript, Mest, in the developing mouse neocortex. Neuroscience 355, 126–140. 10.1016/j.neuroscience.2017.05.003.

Kim, D., Paggi, J.M., Park, C., Bennett, C., and Salzberg, S.L. (2019). Graph-based genome alignment and genotyping with HISAT2 and HISAT-genotype. Nat Biotechnol 37, 907–915. 10.1038/s41587-019-0201-4.

Kirby, S. (2017). Culture and biology in the origins of linguistic structure. Psychon Bull Rev 24, 118–137. 10.3758/s13423-016-1166-7.

Klausberger, T., and Somogyi, P. (2008). Neuronal diversity and temporal dynamics: the unity of hippocampal circuit operations. Science 321, 53–57. 10.1126/science.1149381.

Kobayashi, T., Piao, W., Takamura, T., Kori, H., Miyachi, H., Kitano, S., Iwamoto, Y., Yamada, M., Imayoshi, I., Shioda, S., et al. (2019). Enhanced lysosomal degradation maintains the quiescent state of neural stem cells. Nat Commun 10, 5446. 10.1038/s41467-019-13203-4.

Kolagar, T.A., Farzaneh, M., Nikkar, N., and Khoshnam, S.E. (2020). Human Pluripotent Stem Cells in Neurodegenerative Diseases: Potentials, Advances and Limitations. Curr Stem Cell Res Ther 15, 102–110. 10.2174/1574888X14666190823142911.

Levecque, C., Velayos-Baeza, A., Holloway, Z.G., and Monaco, A.P. (2009). The dyslexia-associated protein KIAA0319 interacts with adaptor protein 2 and follows the classical clathrin-mediated endocytosis pathway. Am J Physiol Cell Physiol 297, C160–168. 10.1152/ajpcell.00630.2008.

Love, M.I., Huber, W., and Anders, S. (2014). Moderated estimation of fold change and dispersion for RNA-seq data with DESeq2. Genome Biol 15, 550. 10.1186/s13059-014-0550-8.

Meng, H., Smith, S.D., Hager, K., Held, M., Liu, J., Olson, R.K., Pennington, B.F., DeFries, J.C., Gelernter, J., O’Reilly-Pol, T., et al. (2005). DCDC2 is associated with reading disability and modulates neuronal development in the brain. Proc Natl Acad Sci U S A 102, 17053–17058. 10.1073/pnas.0508591102.

Molofsky, A.V., and Deneen, B. (2015). Astrocyte development: A Guide for the Perplexed. Glia 63, 1320–1329. 10.1002/glia.22836.

Paracchini, S., Thomas, A., Castro, S., Lai, C., Paramasivam, M., Wang, Y., Keating, B.J., Taylor, J.M., Hacking, D.F., Scerri, T., et al. (2006). The chromosome 6p22 haplotype associated with dyslexia reduces the expression of KIAA0319, a novel gene involved in neuronal migration. Hum Mol Genet 15, 1659–1666. 10.1093/hmg/ddl089.

Pertea, M., Pertea, G.M., Antonescu, C.M., Chang, T.C., Mendell, J.T., and Salzberg, S.L. (2015). StringTie enables improved reconstruction of a transcriptome from RNA-seq reads. Nat Biotechnol 33, 290–295. 10.1038/nbt.3122.

Pletikos, M., Sousa, A.M., Sedmak, G., Meyer, K.A., Zhu, Y., Cheng, F., Li, M., Kawasawa, Y.I., and Sestan, N. (2014). Temporal specification and bilaterality of human neocortical topographic gene expression. Neuron 81, 321–332. 10.1016/j.neuron.2013.11.018.

Powers, N.R., Eicher, J.D., Miller, L.L., Kong, Y., Smith, S.D., Pennington, B.F., Willcutt, E.G., Olson, R.K., Ring, S.M., and Gruen, J.R. (2016). The regulatory element READ1 epistatically influences reading and language, with both deleterious and protective alleles. J Med Genet 53, 163–171. 10.1136/jmedgenet-2015-103418.

Schatschneider, C., and Torgesen, J.K. (2004). Using our current understanding of dyslexia to support early identification and intervention. J Child Neurol 19, 759–765. 10.1177/08830738040190100501.

Shaywitz, S.E., and Shaywitz, B.A. (2005). Dyslexia (specific reading disability). Biol Psychiatry 57, 1301–1309. 10.1016/j.biopsych.2005.01.043.

Stockinger, P., Maitre, J.L., and Heisenberg, C.P. (2011). Defective neuroepithelial cell cohesion affects tangential branchiomotor neuron migration in the zebrafish neural tube. Development 138, 4673–4683. 10.1242/dev.071233.

Szalkowski, C.E., Fiondella, C.G., Galaburda, A.M., Rosen, G.D., Loturco, J.J., and Fitch, R.H. (2012). Neocortical disruption and behavioral impairments in rats following in utero RNAi of candidate dyslexia risk gene Kiaa0319. Int J Dev Neurosci 30, 293–302. 10.1016/j.ijdevneu.2012.01.009.

Velayos-Baeza, A., Toma, C., Paracchini, S., and Monaco, A.P. (2008). The dyslexia-associated gene KIAA0319 encodes highly N-and O-glycosylated plasma membrane and secreted isoforms. Hum Mol Genet 17, 859–871. 10.1093/hmg/ddm358.

Wu, G.D., Li, Z.H., Li, X., Zheng, T., and Zhang, D.K. (2020). microRNA-592 blockade inhibits oxidative stress injury in Alzheimer’s disease astrocytes via the KIAA0319-mediated Keap1/Nrf2/ARE signaling pathway. Exp Neurol 324, 113128. 10.1016/j.expneurol.2019.113128.

Zhang, J., Uchiyama, J., Imami, K., Ishihama, Y., Kageyama, R., and Kobayashi, T. (2021). Novel Roles of Small Extracellular Vesicles in Regulating the Quiescence and Proliferation of Neural Stem Cells. Front Cell Dev Biol 9, 762293. 10.3389/fcell.2021.762293.

Zhao, H., Chen, Y., Zhang, B.P., and Zuo, P.X. (2016). KIAA0319 gene polymorphisms are associated with developmental dyslexia in Chinese Uyghur children. J Hum Genet 61, 745–752. 10.1038/jhg.2016.40.

