## Supplementary figures and images for "Dyslexia associated gene, *KIAA0319*, regulates cell cycle during human neuroepithelium development"

### Supplementary Figures

#### Suppl Figure 1

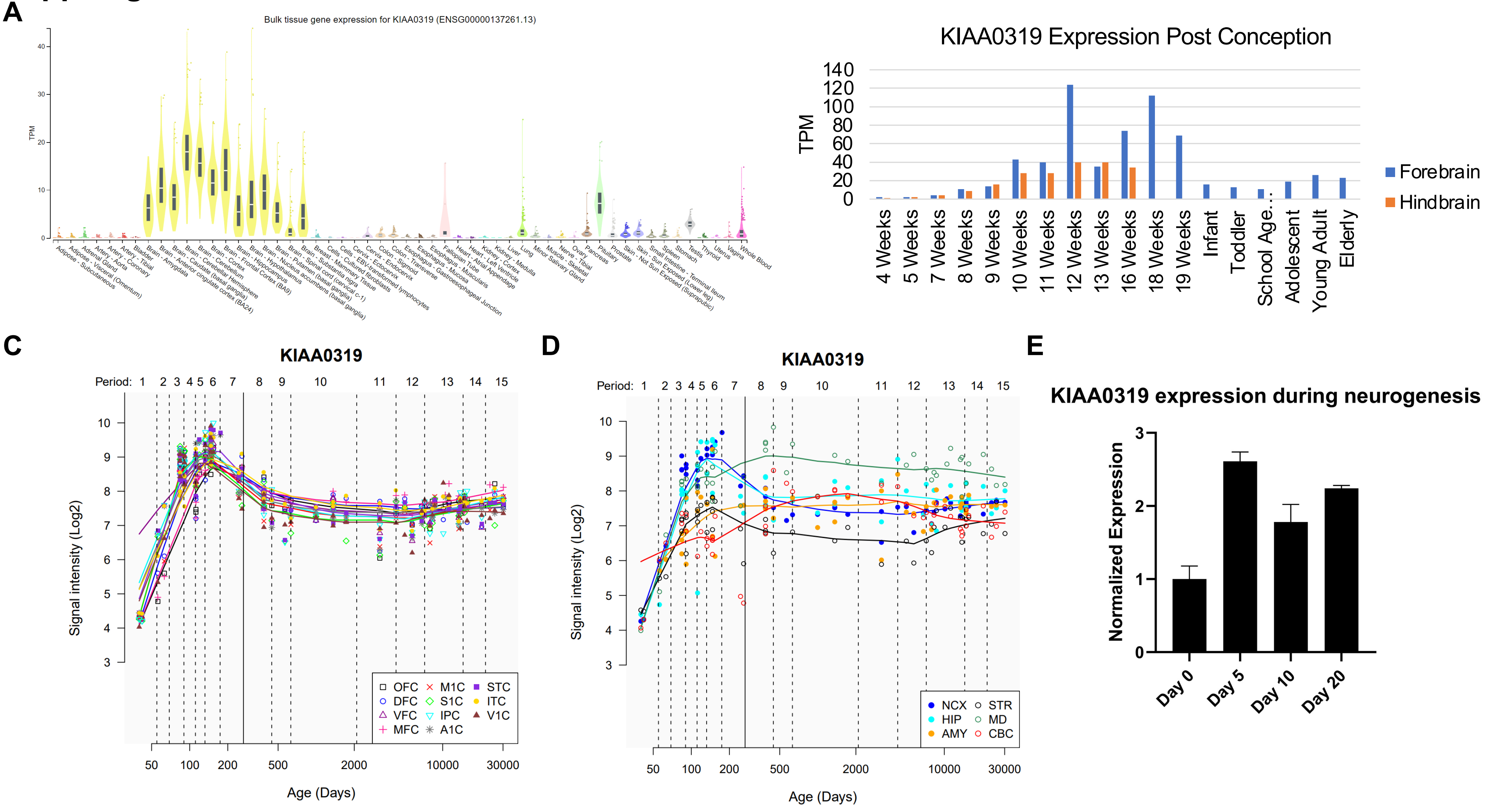

#### Suppl Figure 2

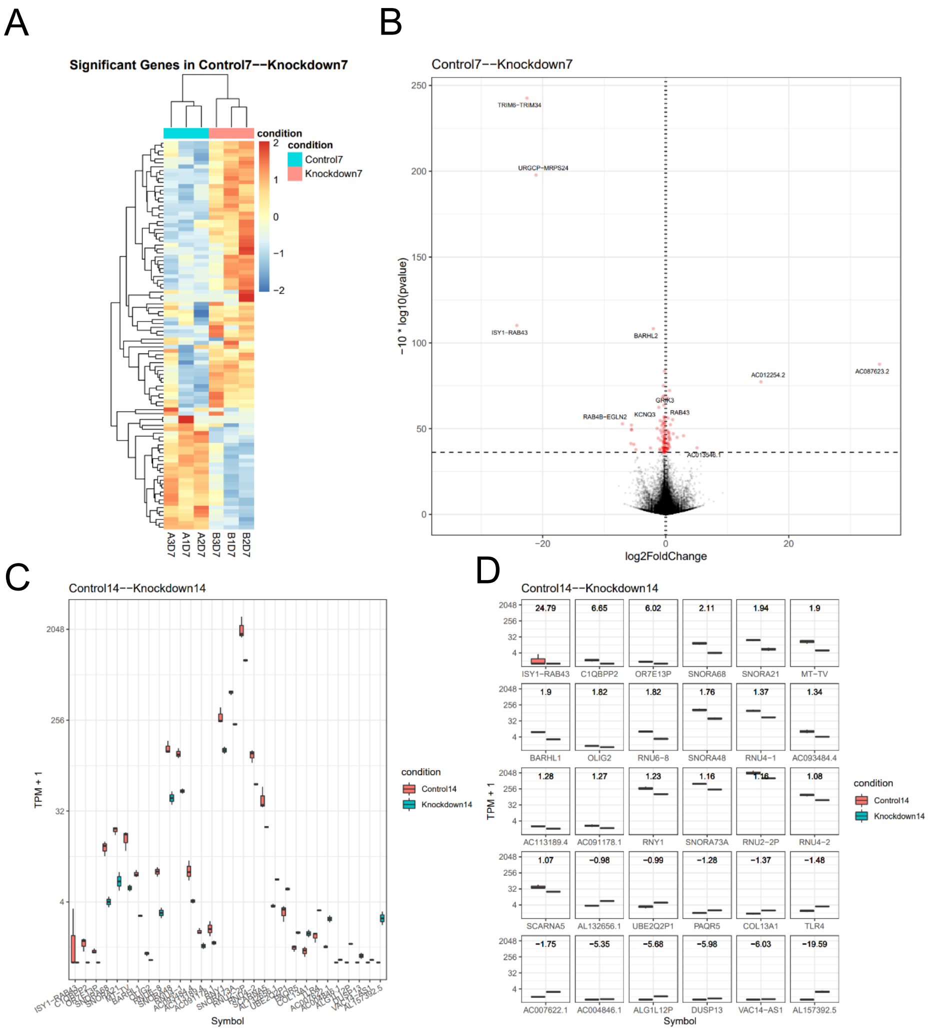

#### Suppl Figure 3

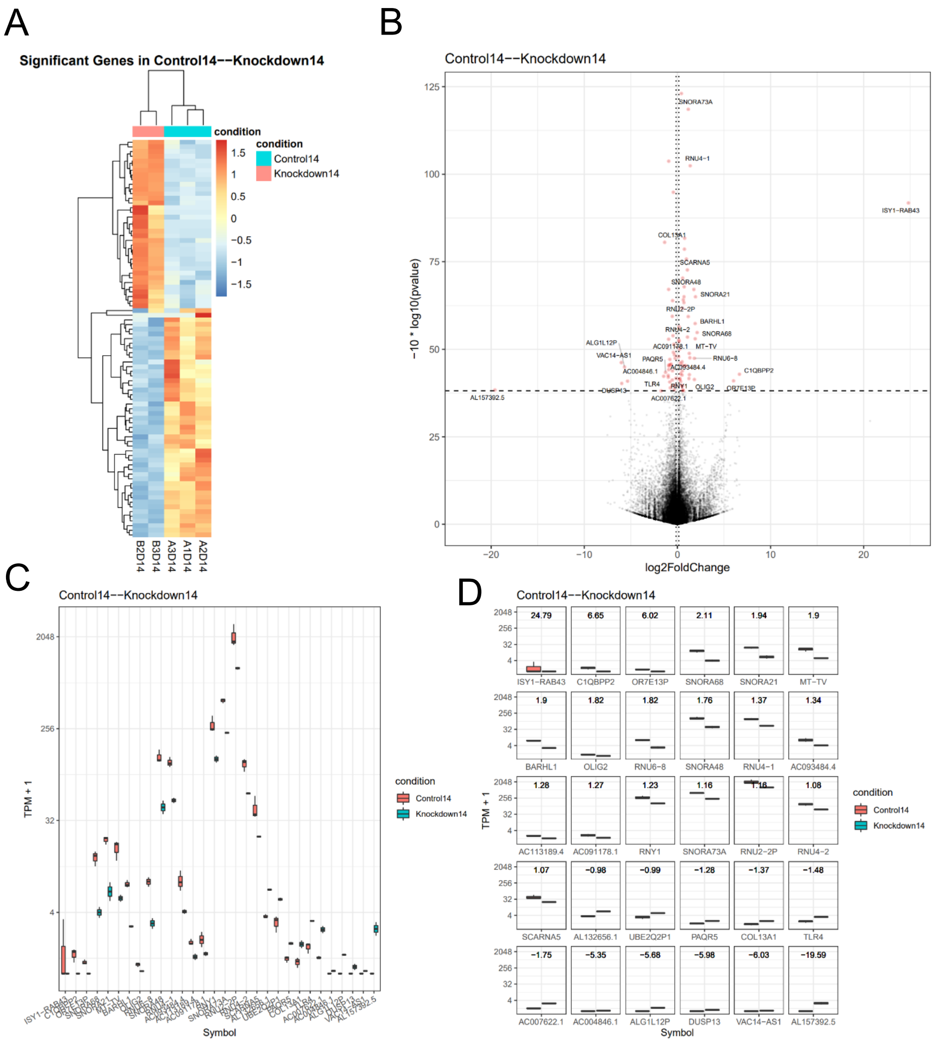

#### Suppl Figure 4

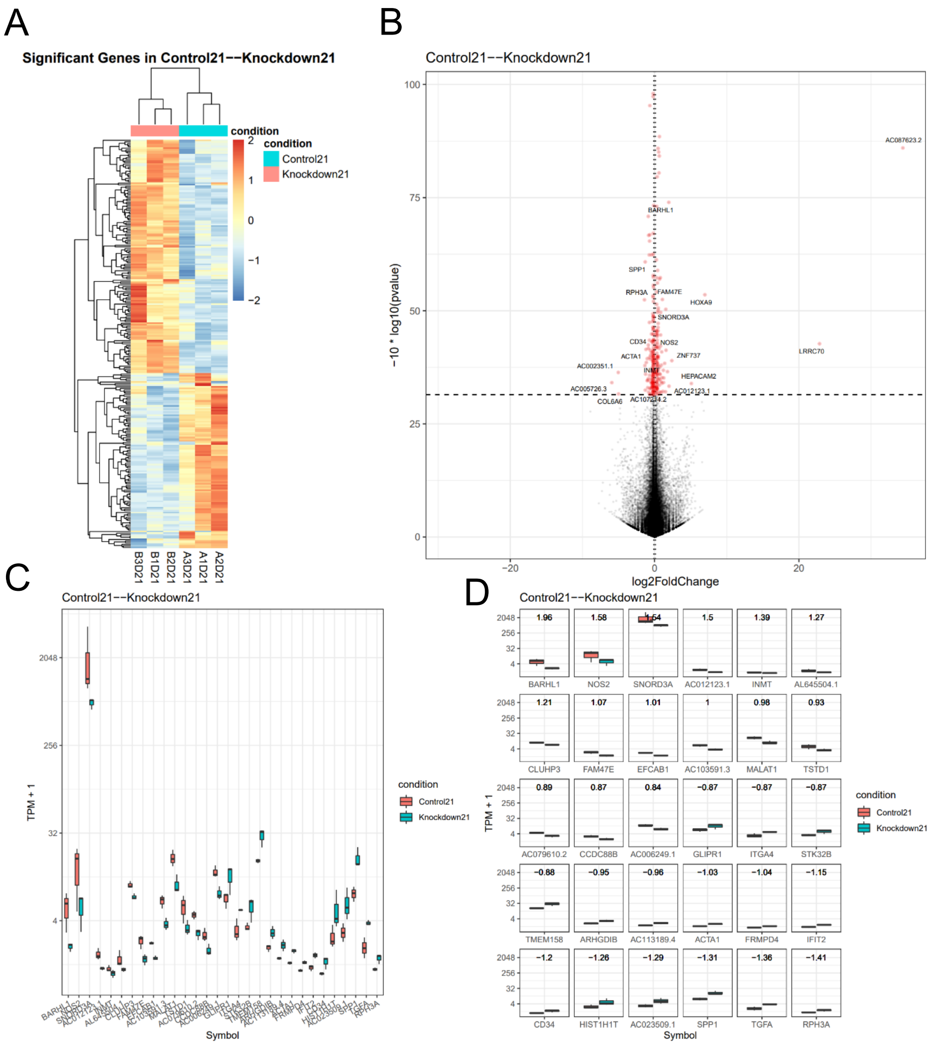

#### Suppl Figure 5

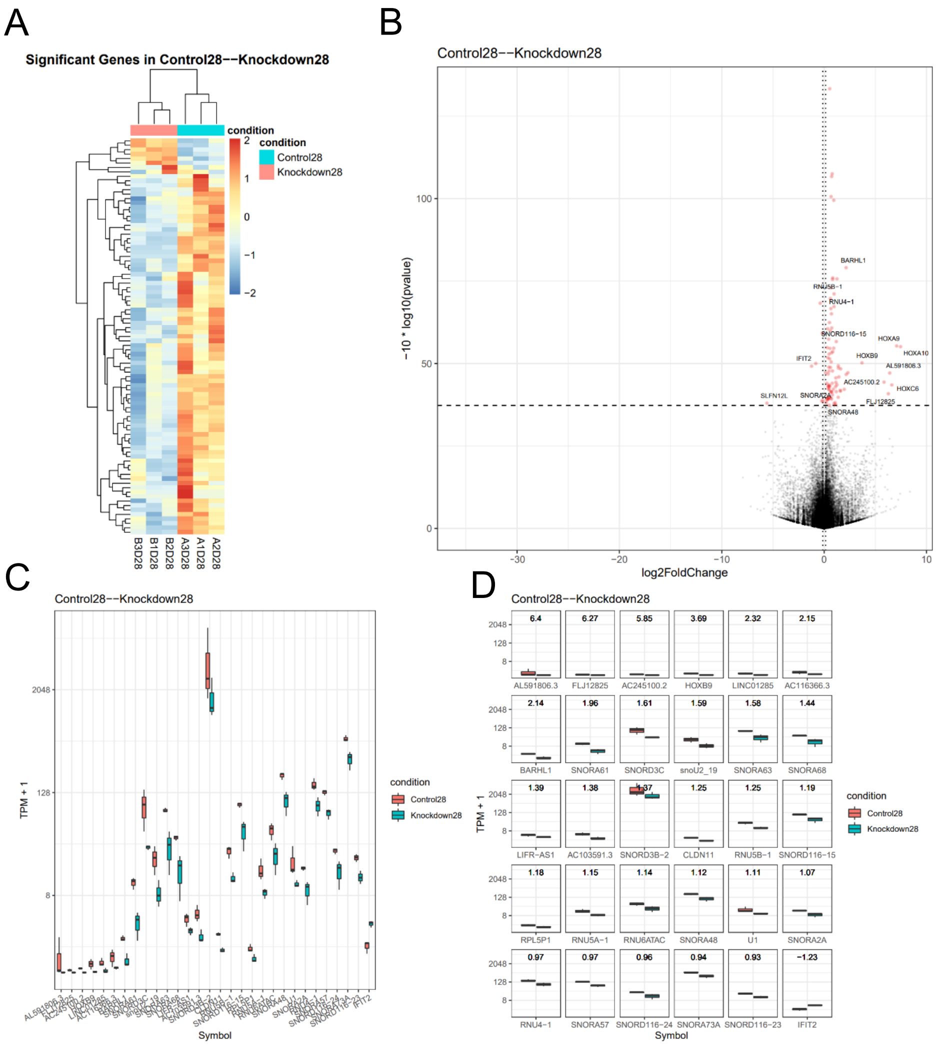
